# Estrogenic Activity in Tampon Products

**DOI:** 10.1101/2025.07.20.665741

**Authors:** E.S. Sutherland, G. Manodara, A. Gillon, K. Stevens, A. Kauff, A.K. Heather

## Abstract

Tampons are widely used menstrual products with prolonged mucosal contact, raising questions about their potential role in exposing females to endocrine-disrupting chemicals (EDCs). This study evaluated estrogenic activity in tampon extracts using a cell-based estrogen receptor reporter bioassay. Synthetic and organic tampons sourced from domestic and international markets—both synthetic and organic—were tested for estrogenic bioactivity. Of the 18 brands analyzed, estrogenic activity was detected in nearly half, independent of material type. Comparative chemical analysis of one estrogenic-active brand versus one non-active brand via high-resolution mass spectrometry identified various plasticizers, surfactants, and fragrance agents, that were present in higher concentrations in the estrogenic-active brand. The list of chemicals included known EDCs, such as phthalates and alkylphenols, that may be responsible for the observed estrogenic activity. Notably, estrogenic activity varied by brand, suggesting formulation-dependent risk. Although the detected in vitro activity levels were low, the findings demonstrate that compounds capable of activating estrogen receptors can leach from tampons. These results highlight the importance of including endocrine bioactivity assays in tampon safety assessments and suggest that safer formulations are achievable. Further investigation is warranted to assess long-term health implications of cumulative low-level EDC exposure from menstrual products.

## 1.0 Introduction

Tampons are widely used menstrual products with reports of up to 85% of females favouring them over other products in Western countries (1,2). In order to manage bleeding, tampons are inserted into the vagina for hours at a time resulting in prolonged contact with the highly vascularized vaginal mucosa. This contact promotes absorption through the walls of the vaginal mucosa bypassing hepatic first-pass metabolism. For certain compounds, such as estradiol, this leads to higher serum concentrations compared to oral administration (2,3). The high usage of tampons together with the intravaginal absorbency has led to recent questioning of the potential of tampons to act as a source of chemical exposure.

Tampons comprise a core absorbent material of cotton, rayon or a blend of viscose/rayon/cotton. Tampons comprised of 100% cotton are marketed as organic products differentiating them from conventional tampons that are inherently more synthetic in composition. The core is contained by an outer layer that can be cotton veil, polyester or polypropylene. The string used for vaginal removal is typically made of cotton, but in the synthetic brands can be a blend of polyester and polypropylene. The string is often bleached and coated with a surfactant to prevent fraying. During manufacture, functional agents can be added to the tampons including plasticizers to make them more flexible, surfactants to assist in fibre processing and fluid absorption, fragrances and deodorants to provide scent, and lubricants and emollients to improve insertion comfort (4).

Tampon exposure in a typical female will on average be for 39 menstrual years equating to 488 menstrual cycles. During this time, she will use approximately 11,700 tampons. A range of chemicals have previously been detected in tampons including dioxins, furans, polycyclic aromatic hydrocarbons, phthalates, parabens, bisphenols, triclocarbon, glyphosate, and volatile organic compounds (5,6). Metals(loids) have also been detected in tampons including lead, arsenic, calcium, zinc amongst others. It is recognised that tampon functionality requires the inclusion of some chemical compounds to provide product integrity, absorbency, flexibility and comfort (4). The presence of chemicals in itself is not inherently harmful; the concern lies in whether these chemicals exist at concentrations that could leach from the tampon material and potentially exhibit hormonal activity-singly, additively or synergistically.

Endocrine-disrupting chemicals (EDCs) adversely impact on the normal functioning of the endocrine system. They can mimic, block or alter the signalling of natural hormones in the body, creating hormonal imbalance. Notably, EDCs can exert their effects at very low concentrations and may have significant effects on human physiology, especially during critical windows of development including fetal life, puberty or pregnancy (7–9). Common examples of EDCs include certain plasticizers such as phthalates, industrial chemicals such as bisphenol A, pesticides like DDT, and personal care product ingredients such as parabens. Many EDCs exert estrogenic bioactivity that alters reproductive function, resulting in reduced ovarian reserve, endometriosis and uterine fibroids, and the disruption of menstrual cycles, as well as an increased risk of breast cancer (10,11).

Cell-based estrogen reporter assays, such as the T47D-KBluc estrogen bioassay (T47D-ER cells), are useful tools for detecting and quantifying estrogenic activity in complex chemical mixtures (12,13). The T47D-R cells have been genetically modified to express a luciferase reporter gene under the control of estrogen-responsive elements (EREs). When the naturally expressed estrogen receptors (ERs) bind estrogenic compounds, they activate the expression of luciferase. The level of luciferase can be quantified as measurable luminescence thereby providing a functional readout of ER activation. The key advantage of this approach is the ability to measure net biological effect of all estrogenic compounds present in a sample. This is especially valuable when assessing the complex mixture of plasticizers, surfactants, emollients, lubricants, and fragrance agents, that may co-exist in tampon material. Many of these may only be weak agonists in isolation, but potentially produce biological effects in combination.

This study aimed to evaluate the estrogenic activity of synthetic and organic tampon extracts using the T47D-KBluc estrogen bioassay. We sought to determine whether differences in chemical content could explain differences in estrogenic bioactivity observed in vitro, and to explore what these findings might suggest about the potential for tampon-derived chemicals to interact with ERs.

## 2. Materials & Methods

### 2.1 Tampon sample selection

Tampon products from multiple manufacturers, brands, product lines, absorbances and counties of origin were selected for study. We tested a total of 18 unique brand-produc-line-absorbency combinations, representing 13 brands, across synthetic and organic compositions (Table 1). We purchased tampons between July 2023 and October 2024 from supermarket stores in New Zealand, and from online retailers (for European and USA purchases). Tampons purchased in New Zealand were not the same products as purchased from the USA, although there was overlap for two brands (Table 1). One brand sent their products directly to the laboratory. We tested two tampons taken at random from each of the 18 products.

**Table 1:** Tampon brands tested in this study.

| Brand | Tampon material | Country of Origin | Size | Estrogen activity detected |
| --- | --- | --- | --- | --- |
| A | Certified organic cotton | New Zealand | Regular | No |
| B | 100% non-chlorine bleached rayon polyester, polyethylene, polyester string | USA | Regular | No |
| C | 100% certified organic cotton, including string. Wrapper made from cellulose. | Europe | Regular | No |
| D | Rayon made from cellulose fibres, polyester, polyethylene, paraffin wax, pigment white 6 (Titanium dioxide) | USA | Regular | Yes |
| E | Top sheet 100% organic cotton, core FSC® certified material, plant-based | Australia | Regular | Yes |
| F | Chlorine-free bleached rayon (derived from plant materials), polypropylene/polyester cover, polyester/cotton string | Australia | Regular | Yes |
| G | 100% certified organic cotton | Europe | Regular | Yes |
| H | 100% certified organic cotton including veil, core, string. Paper wrapper, cardboard box | Europe | Regular | Yes |
| I | Organic cotton (core, string, water repellent wax) | Israel | Super | No |
| J | Organic cotton (core, string, water repellent wax) | Slovenia | Super | No |
| K | Organic cotton (core, string, water repellent wax) | Israel | Regular | No |
| L | Organic cotton (core, string, water repellent wax) | Israel | Super | No |
| M | Organic cotton | Europe | Regular | No |
| N | Rayon, cotton fiber, polyester, polysorbate 20, wax blend (paraffin, butyl stearate, carbauba wax, polymer wax dispersion) | USA | Regular | Yes |
| O | 100% cotton core | USA | Regular | No |
| P | 100% cotton core | USA | Regular | Yes |
| Q | Rayon, water, polyester, polyethylene, paraffin wax, pigment white 6 (titanium dioxide) | USA | Regular | Yes |
| R | Rayon, polyester, purified cotton | USA | Regular | No |

**Table 2:** Orbitrap MS parameters.

| Parameter | Setting |
| --- | --- |
| Positive Ion (V) | 3400 |
| Negative Ion (V) | 2000 |
| Sheath Gas (Arb) | 5 |
| Sweep Gas (Arb) | 5 |
| ITT (°C) | 350 |
| Vaporiser (°C) | 400 |
| AGC Target | Standard |

### 2.2 Sample preparation and analysis

Each tampon was weighed and two 1 g portions were cut from each tampon. These were soaked in 10 mL ethanol:milliQ water (50% v/v) for 4 hours then agitated by 10 min sonication, 40 min shaking (200-210 rpm), 10 min sonication and then vortexing for 2 mins. The exudate was centrifuged for 15 mins at 4000g. Five mL of the supernatant was then loaded into a pre-washed SPE column (2 g Bond Elut, Agilent Technologies) that was pre-conditioned with 3 mL methanol, followed by 3 mL sodium acetate (pH 4.8). The column was washed sequentially with 2 mL laboratory grade water, 1.5 mL sodium carbonate (10% w/v), 2 mL laboratory grade water and 2 mL laboratory grade water/methanol (1:1 v/v). After washing, the column was air dried before the sample was eluted with 4 mL acetonitrile. The solvent was dried and sample extract reconstituted in 200 μL 100% ethanol (herein “samples”). For testing, 2 μL of the sample was added to 198 μL DMEM F-12 phenol-red free medium. From this dilution, 100 μL was mixed with 100 μL DMEM F-12 phenol-red free medium. This dilution series was continued for 2 more dilutions and all four dilutions were tested.

### 2.3 Estrogen receptor bioassay

T47D-KBluc reporter cells (ATCC CRL-2865) were grown to 90% confluence in a 550 mL cell culture flask with RPMI 164 medium supplemented with 10% (v/v) Hyclone fetal bovine serum (GE Lifesciences) and Geneticin (75 μgmL^-1^, G148, Life Technologies). Cells were then seeded at 5X10^4^ cells per 100 μL DMEM F-12 phenol-red free medium per well of a 96-well culture plate. The medium was supplemented with charcoal-stripped FBS (10% v/v) and G148 (75 μgmL^-1^). Cells were then incubate at 37°C for 24 hours under 5% CO_2_.

For treatment with estradiol standards or samples, medium was removed from the cells by aspiration, and replaced with 90 μL of fresh DMEM F-12 phenol-red free medium, supplemented with charcoal-stripped FBS (2% v/v) and G148 (75 μgmL^-1^). 10 μL of the estradiol standards or the sample dilutions were added to 90 μL medium and the cells were incubated at 37°C for 24 hours under 5% CO_2_. After extraction and sample dilution, the highest sample concentration represented a 60-fold dilution of the original extract (25-fold concentration by extraction with a 1 in 1500 dilution in cell culture medium the time of cell exposure).

For estradiol standards preparation, a 2 μM stock solution of estradiol in 100% ethanol was diluted as 15 μL in 285 μL of DMEM F-12 phenol-red free medium. From this 100 nM stock, 100 μL was added to 200 μL DMEM F-12 phenol-red free medium. This 1:3 dilution was continued for 10 more dilutions and all 11 dilutions were used to generate the estradiol dose response curve.

For all assay runs, a solvent control was analysed. This was prepared by adding 2 μL of 100% ethanol to 298 μL DMEM F-12 phenol-red free medium. From this dilution, 100 μL was added to 200 μL DMEM F-12 phenol-red free medium.

After incubation, media was aspirated from all wells and 100 μL BrightGlo substrate (Promega) was added. After a 2 min incubation at room temperature, the BrightGlo was transferred to a white opaque 96-well plate and luminescence reported using a SpectraMax i3x plate reader (Molecular Devices).

### 2.4 Liquid Chromatography – High Resolution Mass Spectrometry (LC-HRMS) protocol

Extraction of potential EDCs from tampons was performed as above. For analysis, the 200 μL extract from 7 tampons of Brand A or D were combined in order to have the extract volume needed for analysis. Separation was performed with an Ultimate 3000 HPLC coupled to a Thermo Exploris 480 Orbitrap Mass Spectrometer. LC separation was performed on an Acquity UPLC BEH C18 column (100 x 2.1 mm ID, 1.7 μm, Waters) at 55°C in a linear gradient (20-95% over 29 minutes) with water and methanol as the mobile phases, both containing 0.1% formic acid. The mass spectrometer was equipped with an electron spray ionization source (H-ESI) operated in positive and negative mode with parameters described in Table 1. Data were acquired over the mass range of 100-1000 Da using full scan MS mode at 120,000 resolution in conjunction with an internal mass calibration (EASY-IC). This allowed a non-targeted approach to the analysis. In addition, data dependent mass scan (ddMS2) was used to collect fragmentation data at 15,000 resolution at 30 and 80V HCD collision energy. Sensitivity of likely target analytes were checked to ensure optimal ionisation and separation parameters.

Data generated from the LC-HRMS were processed using Compound Discoverer 3.3 SP2 application (Thermo Scientific). Retention times (RT) were automatically aligned, and peak assignment was performed with a mass tolerance of 5 ppm and RT tolerance of 0.1 min. The workflow searched for multiple adducts as well as isotope pattern detection (Br;Cl). Identification was done by a local database and other data sources such as MassList, Chemspider and mzCloud.

### 2.5 Analysis

Luminescence values were first corrected by subtracting the mean signal from blank wells (media and ethanol only). Estradiol was used as a positive control to generate a standard curve using a four-parameter logistic regression to fit the data (0.001-100 nM) (PRISM Graphpad v10). For samples, each tampon was tested as duplicate 1 g subsamples. Each subsample was assayed in duplicate for each of the four dilutions tested. Subsamples were averaged to provide the mean per tampon. Statistical significance between treatments was assessed using one-way ANOVA followed by Tukey’s post-hoc test, with p-values < 0.05 considered significant. The limit of detection (LOD) was defined as the mean of the vehicle control plus three times the standard deviation. Samples eliciting responses above the LOD were classified estrogenically active. All analyses were performed using Graphpad PRISM v10.4.1.

## 3.0 Results

### 3.1 Characterization of the T47D-luc estrogen bioassay

The sensitivity of the T47D-KBluc estrogen bioassay was evaluated to determine its suitability for testing extracts derived from tampons. Figure 1 shows a clear sigmoidal dose-response curve for E2. The calculated EC_50_ for T47D-luc cells under our experimental conditions was approximately 14.5 pM. The bioassay demonstrated a limit of detection at approximately 0.5 pM in keeping with a previous report (14).

**Figure 1:**
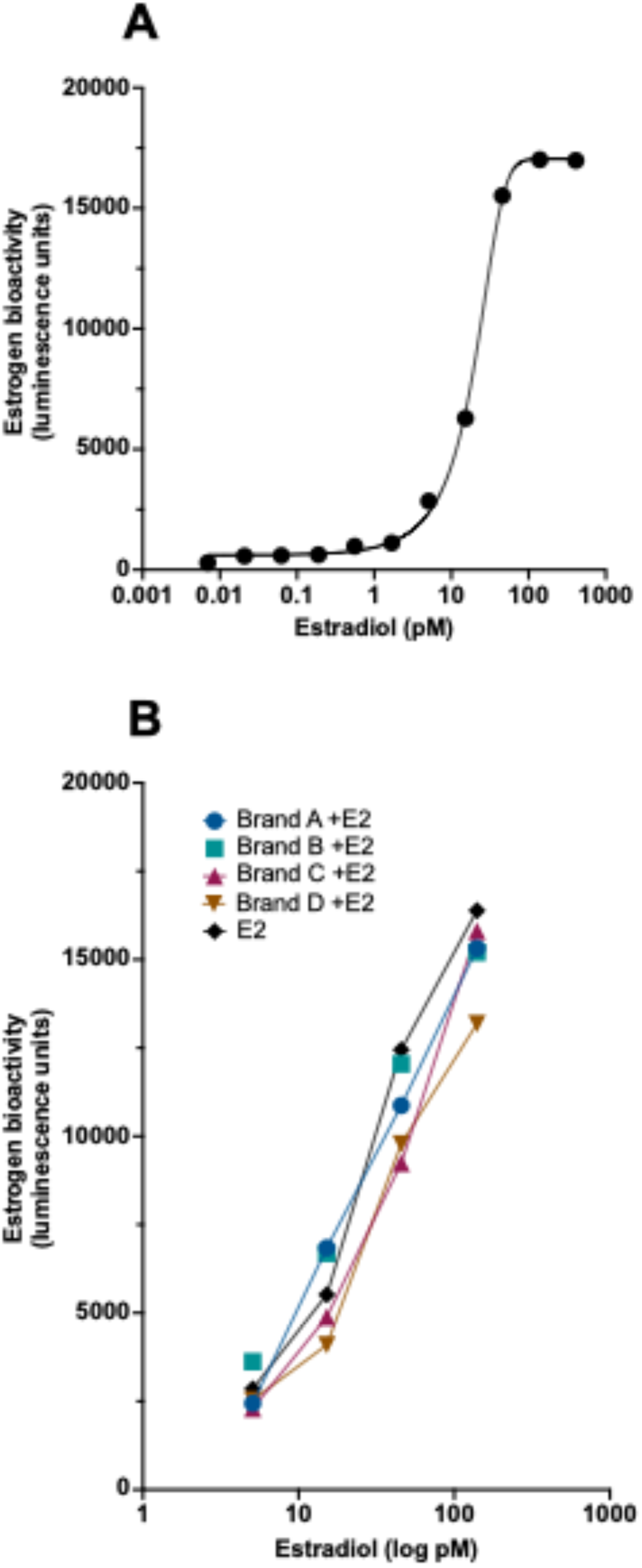
Sensitivity of the T47D-ER luciferase assay and validation of estradiol recovery from tampon extracts. A. Dose-response curve of T47D-ER cells treated with increasing concentrations of estradiol (E2). Estrogen bioactivity is shown as luminescence (arbitrary units). B. Estrogen bioactivity in T47D-ER cells treated with E2 (2 nM) spiked into extracts from four tampon brands (A-D), compared to E2 alone.

### 3.2 Estrogenic activity measured in E2-spiked tampons

In order to demonstrate that the T47D-KBluc could detect estrogenic activity from tampon extracts, four tampons from four different brands were spiked with estradiol at a concentration of 2 nM. Each tampon was then soaked in a 1:1 water:ethanol solution for 4 hours, mimicking the recommended duration of tampon use in females. Post-soaking, the spiked tampons underwent the sonication, extraction and dilution protocol before being tested for estrogenic activity using the T47D-KBluc assay.

Figure 2 illustrates the estrogenic activity observed from four tampon brands spiked with estradiol. The results demonstrate that the assay is able to accurately detect estrogenic activity from the tampon exudates post-extraction and dilution. Detected estrogenic activity levels closely resembled those of estradiol alone. Specifically, at the highest dilution tested, estrogen recapture efficiencies were determined to be 93.5% for Brand A, 92.8% for Brand B, 94.3% for Brand C, and 80.5% for Brand D.

**Figure 2:**
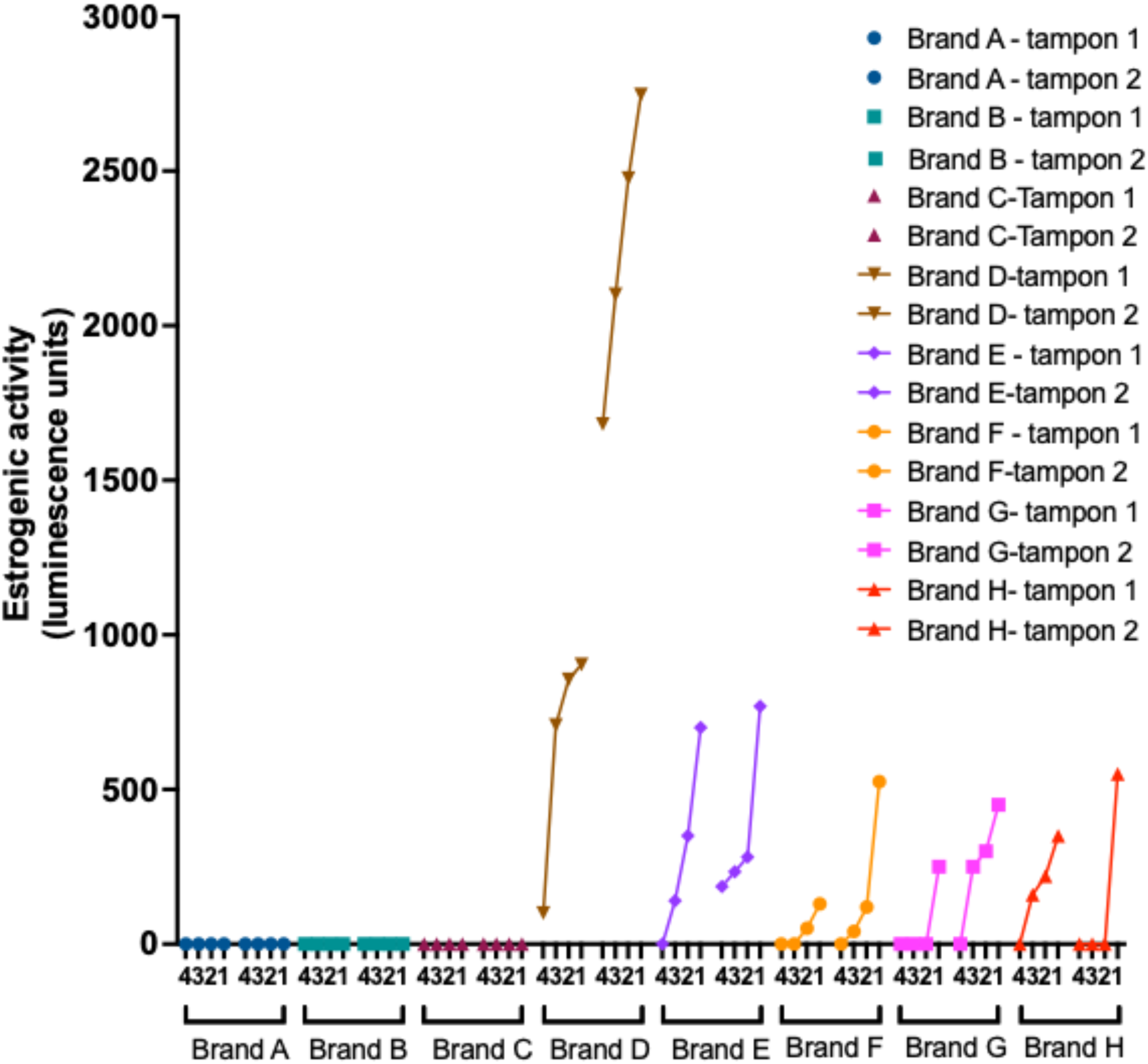
Estrogenic activity in multiple brands of tampon extracts. Estrogenic activity measured in T47D-ER cells treated with four serial dilutions (x-axis 4321, 1=most concentrated, 4=most diluted) of extracts from eight tampon brands (A-H). Luminescence values represent ER-mediated bioactivity (estrogenic activity, arbitrary units). Data points represent the mean of replicates, from two tampons.

### 3.3 Estrogenic activity measured in tampon extracts sourced from New Zealand supermarkets

We next evaluated estrogenic activity from tampon extracts across eight brands sourced from supermarkets in New Zealand, except for Brand A, which was provided directly by the manufacturer (Table 1). Two tampons per tampon packet were tested. The results presented in Figure 3 indicate that extracts from Brands A, B, and C exhibited no detectable estrogenic activity indicating that both tampons from these brands do not leach estrogen-mimicking compounds under the experimental conditions. In contrast, Brands D through H demonstrated clear estrogenic bioactivity, the level of which varied across brands and predictably decreased with increasing dilution (dilution 1 representing the highest concentration and dilution 4 the most dilute). The two tampons from each positive brand showed variability likely reflecting the differences in concentration of leached chemicals. Although the overall magnitude of estrogenic activity was relatively low—falling toward the lower end of the estradiol standard response curve (as shown in Figure 1)—the clear dose-dependent reduction observed in each dilution series confirms the presence of genuine estrogenic activity, as opposed to random signal fluctuations or background noise. The experimental setup resulted in a 60-fold dilution of the sample at cell exposure indicating that actual estrogenic activity present in the leachate is likely to be significantly higher than the unadjusted values shown.

**Figure 3:**
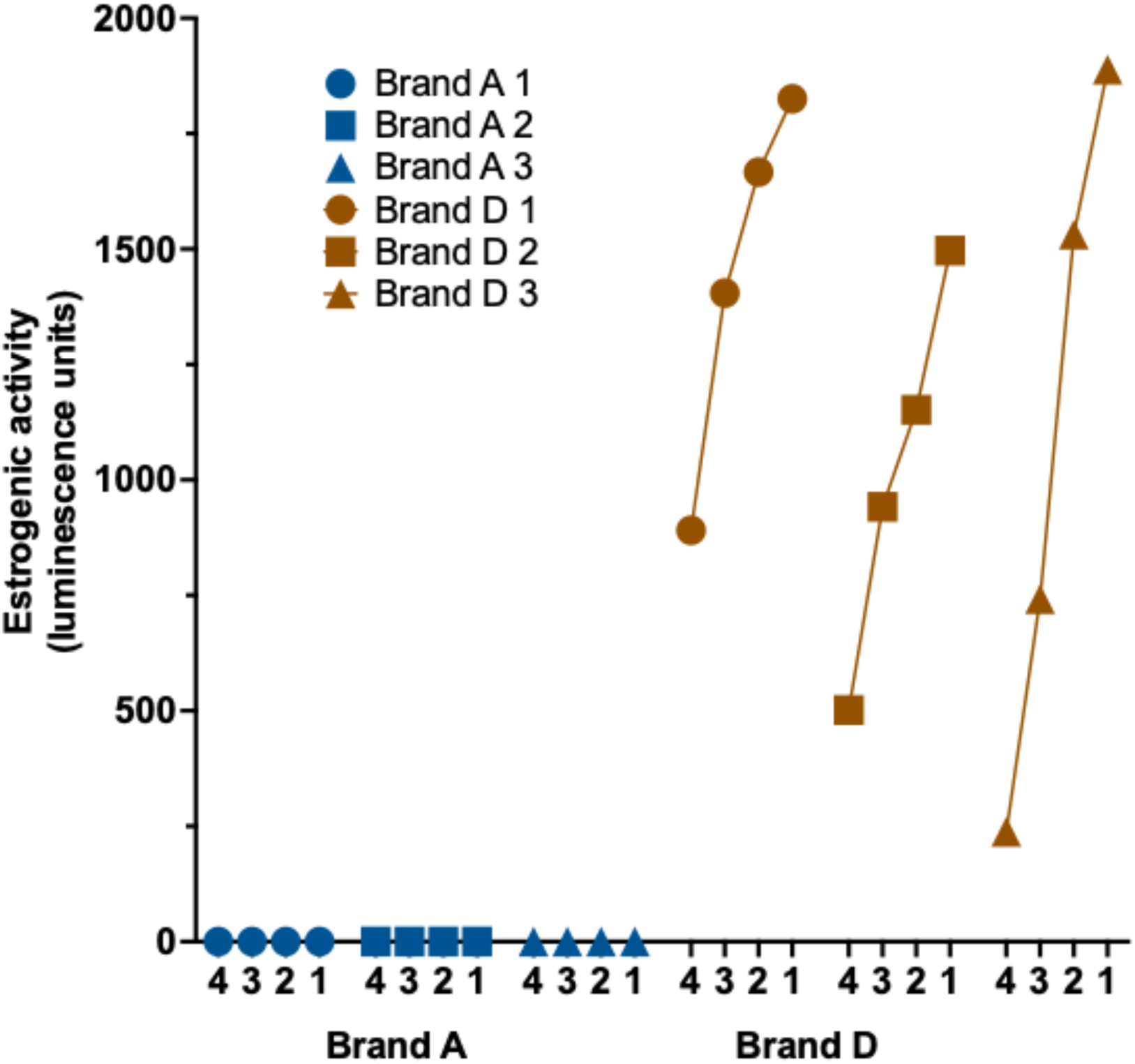
Estrogenic activity in multiple batches of tampon extracts from two brands. Estrogenic activity measured in T47D-ER cells treated with serial dilutions (x-axis 4321, 1=most concentrated, 4=most diluted) of extracts from 3 independent batches (1–3) of Brand A and Brand D. Data points represent the mean of replicates, from two tampons per batch.

### 3.4 Testing batches of two selected tampon brands for estrogenic activity

We next analyzed three separate batches of two selected brands: one determined to lack estrogenic activity (Brand A) and one positive for estrogenic activity (Brand D). For Brand A, no estrogenic activity was detected across all three batches, supporting the conclusion that Brand A does not contain detectable levels of estrogenic compounds under the experimental conditions tested (Figure 4). Conversely, all three batches of Brand D exhibited estrogenic activity, albeit with notable intra-batch variability. This observed variability for batches most likely reflects the rate of chemicals leaching into the exudate because of concentration differences in the tampons and/or variation in the recovery efficiency during SPE extraction.

**Figure 4:**
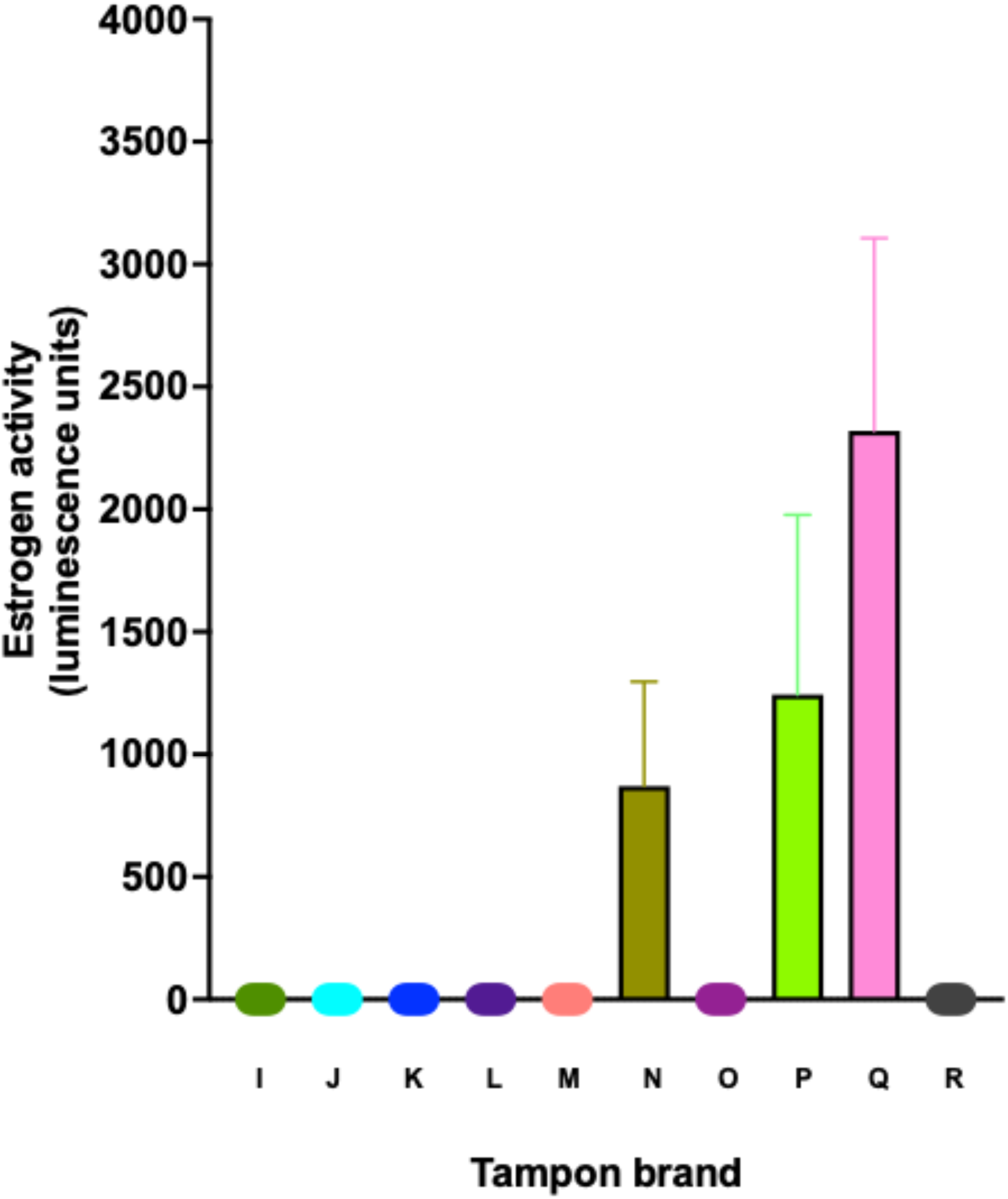
Estrogenic activity of internationally-sourced tampon extracts. Estrogenic activity measured in T47D-ER cells treated with extracts from tampons sourced from 9 international brands (I-R). Bioactivity is reported as mean luminescence units (mean +/- SEM) from two tampons each with 4 replicates.

### 3.5 Estrogenic activity in internationally-sourced tampon brands

The data obtained thus far were from tampons sourced from New Zealand supermarkets. We next tested brands that were purchased internationally. For each brand, 2 tampons were tested, and the data shows that Brands I to M, O and R showed no measurable estrogenic activity. Brand N, P and Q showed measurable estrogenic activity.

### 3.6 Estrogenic activity in tampon brands with and without sonication

To help release any potential estrogenic chemicals into the tampon exudate our processing of the tampons involved sonication after soaking for 4 hours. Sonication itself did not cause destruction of the tampon with the core remaining intact, however, it is a step that does not reflect the real-time use of tampons. To determine if sonication artificially created an estrogenic profile for the tampon exudates that would not physiologically occur, we compared the estrogenic activity of Brand A and Brand D after processing with and without a sonication step.

Brand A showed no estrogenic activity with either active-, or absent sonication treatment (Figure 5). Brand D continued to exhibit estrogenic activity with no sonication, although this process did increase estrogenic activity suggesting agitation does promote greater chemical leaching (Figure 5). Together, the results show that passive leaching occurs with forced agitation simply accelerating the process.

**Figure 5:**
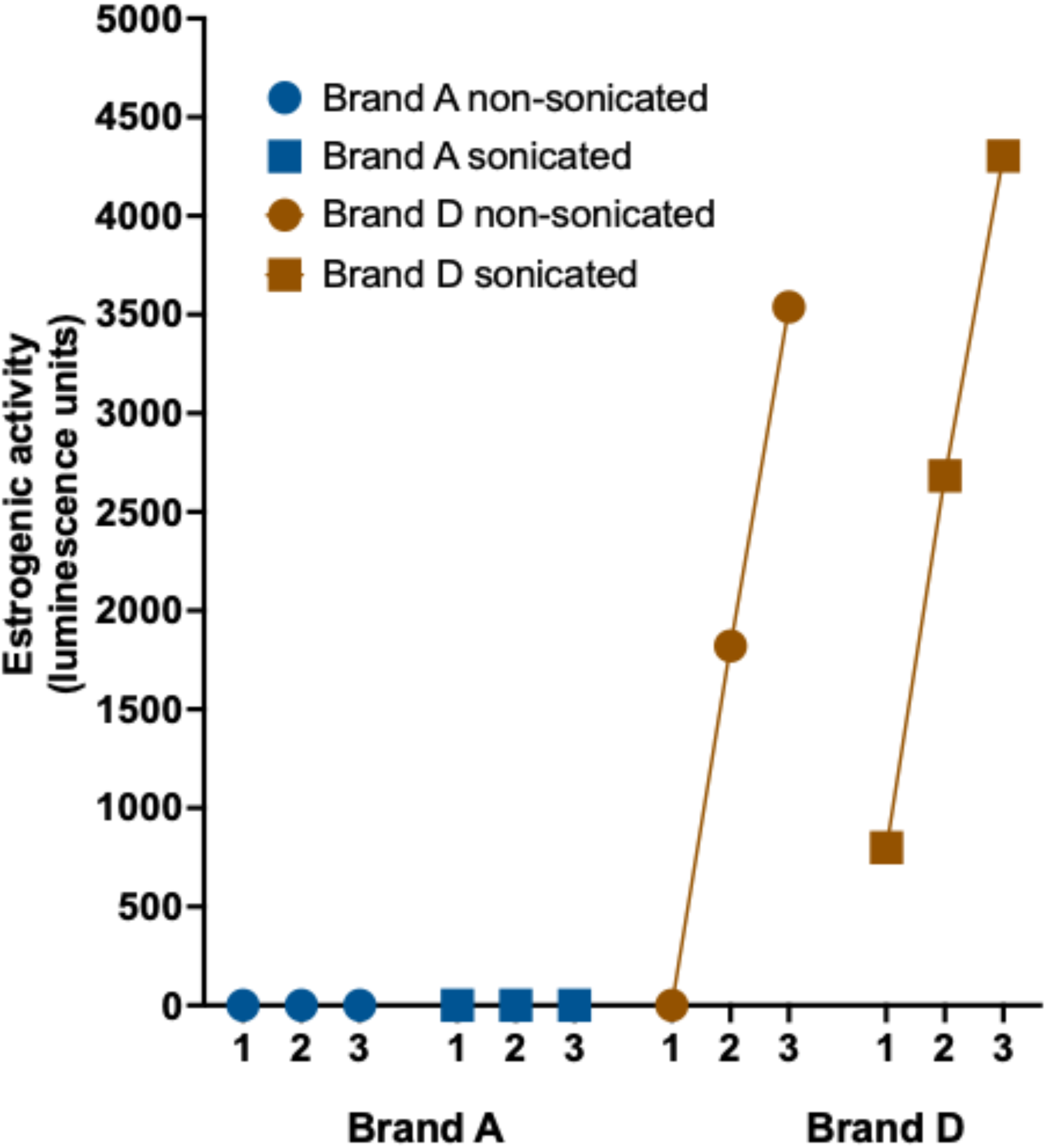
Effect of sonication on estrogenic activity of tampon extracts. Estrogenic activity measured in T47D-ER cells treated with 3 dilutions (X-axis 123, 1=most concentrated, 3= most diluted) of extracts from Brand A and Brand D tampons, prepared with and without sonication. Bioactivity is shown as mean luminescence units from two tampons, each with 4 replicates.

### 3.7 Identification of estrogenic chemicals in tampons exhibiting positive estrogenic activity

Chemical analyses of tampon extract injections (samples A and D) were compared to a blank (ethanol) using LC-HRMS. Thousands of unique chemical species were detected, but only well-defined peaks (rating ≥6.5) and those not present in blanks were retained. Tentative compound identities were assigned using matches to exact mass and isotope patterns, and the most abundant were confirmed through MS2 fragmentation patterns. Only features with strong database matches (FISh score >40) were considered reliable.

Statistical evaluation highlighted significant differences in chemical profiles between extracts, with log2 fold changes >5 used to identify compounds that were strongly enriched in one sample. Peak areas and tentative identities are listed in Table 3, providing a comparative view between samples, although direct comparison between different compounds is limited due to variability in ionization efficiency. Table 3 shows that Brand D exhibited a broader spectrum of identified chemicals, with many detected at levels greater than 1.0E^7^ M (well above the blank background) and measured in Brand A.

**Table 3:**
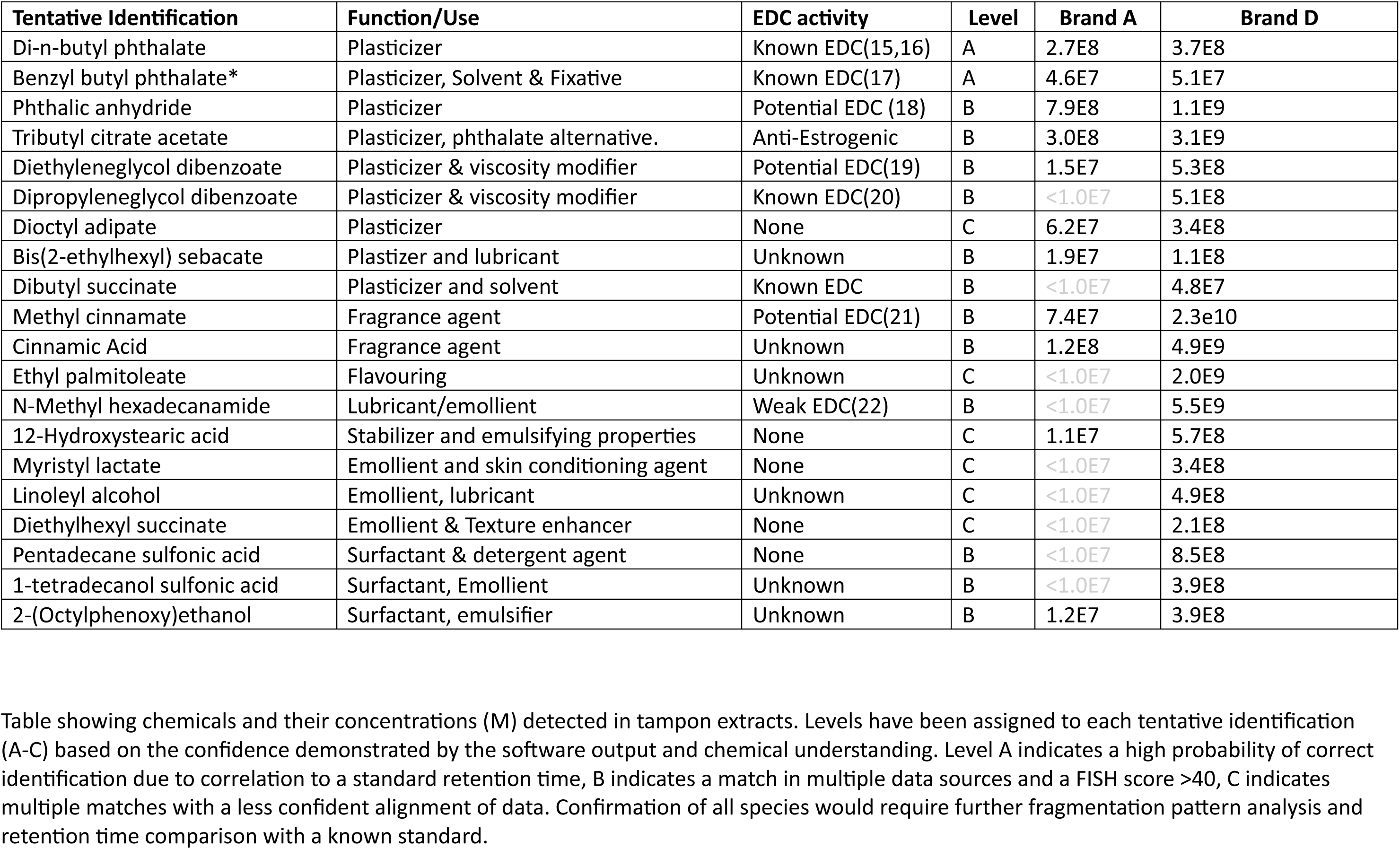
Chemicals identified in tampon extracts by HRMS- Brand A vs Brand D.

## 4.0 Discussion

This study has demonstrated that 8 of 18 tampon brands harbor estrogenic compounds that leach out of the material when soaked over a 4-hour period. The positive tampons included both synthetic and organic brands. Detailed mass spectrometry analysis showed that one positive tampon brand had higher concentrations of known or suspected estrogenic EDCs, including phthalates, 4-tert-octylphenol (OP), 2-octophenoxy ethanol (OPE), hydroxy-1,4-benzoquinone, dioctyl adipate (DEHA) and other plasticizers, fragrances, and surfactant derivatives. These EDCs are very likely responsible for the estrogenic bioactivity of certain tampon brands. We also note that all four batches of Tampon D assayed exhibited estrogenic activity.

The exudates from 18 tampons were tested for estrogenic bioactivity using the well-characterised T47D-KBluc cell-based bioassay (12,13). This cell line represents breast cancer cells that are exquisitely sensitive to estrogenic compounds. The cells were used to test for estrogenic activity in the extracted exudate from tampons, after they had been sonicated and soaked in an ethanolic solution for 4 hours. To ensure the tampon processing steps were suitable for extracting estrogenic compounds, we tested estradiol-spiked tampons and showed for 3 of four brands that subsequent estrogenic bioactivity was > 92% of that expected for the spiked concentration of E2. For the fourth brand, Brand D, E2 recapture efficiency was 80.5%, however it was later shown that this tampon brand harbored estrogenic EDCs and therefore the decrease in E2-induced activity was likely competitively inhibited by leached estrogenic compounds that were now also present in the extracted exudate.

The nature of cell-based estrogenic bioassays is that they measure net effect of all the estrogenic compounds that are present in a sample. This allows the measurement of estrogenic activity from a complex mixture, including additive and synergistic effects. Comparing the percentage of ER bioactivity, and allowing for the 25-fold concentration of the tampon exudate during the extraction process, the E2 equivalence for the average measured estrogenic activity from the tampons is approximately 62.5 pM (17 ng/L). This most likely represents the summed estrogenic activity from several chemicals, rather than the single effect of just one chemical. Individually, these chemicals may not exert strong estrogenic effects, but collectively reach the threshold needed to activate ER-mediated pathways. This likely explains why Brand A was negative for estrogenic bioactivity despite the detection of a small number of potential EDCs in the HRMS analysis. In contrast, Brand D exhibited a wide variety of known and suspected EDCs with several present at over 1.0E^9^ M.

The estrogenic bioactivity measured in the bioassay can be explained by the presence of several known estrogenic EDCs. Phthalates and various plastic additives, surfactants, fragrance and scent-masking agents, as well as lubricant/emollients were all chemically identified using HRMS. The presence of phthalates in tampons is in keeping with a previous study that specifically targeted phthalate detection (23). Phthalates are known for their weak estrogenic effects and strong anti-androgenic effects. Common phthalates, including butyl benzyl phthalate (BBP) and di-n-butyl phthalate (DBP) detected in this study, induced breast cell proliferation, a hallmark of ER agonism at high concentrations (24). The same chemicals can also interfere with aromatase and other steroidogenic enzymes in vitro potentially altering estrogen synthesis and therefore impacting on blood E2 levels (25). These in vitro effects are echoed in human studies, where phthalate exposure has been linked to hypoestrogenism, diminished ovarian reserve, increased risk of premature ovarian failure (26,27), and uterine fibrosis (28–30). With the growing awareness of health risks associated with phthalates, manufacturers have been switching to plasticizers with potentially more favorable profiles. Alternative plasticizers found in this study include diethyl succinate, isophorane, tributyl citrate acetate, dibutyl sebacate, diethylene glycol dibenzoate, dipropylene glycol dibenzoate, tributyl phosphate (TBP), dioctyl adipate (DEHA), bis(2-ethylhexyl) sebacate (DEHS), dimethyl sebacate, and dibutyl succinate. Of these, DEHA at high concentrations was reported to increase breast cell proliferation, in similar fashion to phthalates (31), and DEHS and TBP are both characterized as anti-androgenic, suggesting they may be weakly estrogenic (32). Cinnamate compounds also have weak estrogenic bioactivity due to their structural similarity to endogenous estrogens (21). Octylphenol derivatives were also present in the tampon extracts and these are known to exhibit strong estrogenic activity, even at low concentrations (33). In vivo, octyphenol exposure induces vaginal mucification and histological changes in ovarectomized rats (33).

We provide here the first evidence that compounds leaching from some brands of tampons have demonstrable estrogenic activity using a human ESR1/2-expressing breast cancer cell line. The estrogen equivalence of 62.5 pM or 17 ng/L leached from each tampon is within the range of concern for chronic exposure. Such concentrations can trigger low-dose, non-monotonic effects as has been reported for a number of estrogenic EDCs (34,35). Some of the more studied EDCs have been shown to have the potential to bioaccumulate in tissues, raising the possibility of cumulative exposure over time (36,37). However, at present, the impact of estrogenic EDC-leaching tampons on human physiology or pathophysiology is not known.

The encouraging finding from the study is that tampons can be manufactured free of detectable estrogenic activity. The lack of estrogenic activity in extracts from some organic and synthetic tampons supports a brand-dependent chemical exposure model. While tampon manufacture requires the use of processing chemicals, finishing agents, and packaging that all can harbor estrogenic EDCs, several brands showed no estrogenic activity which highlights that reducing the complexity of chemicals present and reducing individual chemical concentration may be key to achieving safer, non-estrogenic products. The findings raise important questions about regulatory oversight of menstrual products, particularly regarding the use of plasticizers, surfactants, fragrance ingredients known or suspected to be hormonally active. There is a need for labelling requirements for such components in tampons, and the potential need for product safety testing that considers endocrine endpoints, not just irritation and infection risk. The clear difference in estrogenic bioactivity between the different brands reported here suggests that safer formulation is achievable.

In summary, estrogenic activity was demonstrated in tampon extracts in 8 tampon extracts, representing 44% of those tested, in a brand-specific manner. As multiple known and likely unidentified estrogenic compounds exist in tampons, it will be important to use estrogen receptor bioassays of the type reported here to assess and monitor tampon safety in the future. While testing for one individual EDC with standard mass spectrometry approaches is useful, it will not provide a full account of potential estrogenic leaching. Encouragingly, the absence of estrogenic bioactivity in some brands shows that a safer formulation is possible. While it is presently difficult to translate the impact of the in vitro observations to human health, the prospect of bioaccumulating EDCs leaching from tampons into a highly absorbent vagina over a female’s reproductive lifespan, deserves further attention and investigation.

## Conflict of Interest Statement

Heather, Sutherland, Manodara, Gillon, and Kauff were employees of InsituGen Ltd (start-up company) at the time this study was conducted. Heather is the Chief Scientific Officer for InsituGen, and has shares in the company.

The company, Organic Initiative Ltd (New Zealand), provided tampon samples for paid testing but had no input into study design, data analysis, and manuscript preparation.

